# Squeegee: de-novo identification of reagent and laboratory induced microbial contaminants in low biomass microbiomes

**DOI:** 10.1101/2021.05.06.442815

**Authors:** Yunxi Liu, R. A. Leo Elworth, Michael D. Jochum, Kjersti M. Aagaard, Todd J. Treangen

## Abstract

Computational analysis of host-associated microbiomes has opened the door to numerous discoveries relevant to human health and disease. However, contaminant sequences in metagenomic samples can potentially impact the interpretation of findings reported in microbiome studies, especially in low biomass environments. Our hypothesis is that contamination from DNA extraction kits or sampling lab environments will leave taxonomic “bread crumbs” across multiple distinct sample types, allowing for the detection of microbial contaminants when negative controls are unavailable. To test this hypothesis we implemented Squeegee, a *de novo* contamination detection tool. We tested Squeegee on simulated and real low biomass metagenomic datasets. On the low biomass samples, we compared Squeegee predictions to experimental negative control data and show that Squeegee accurately recovers known contaminants. We also analyzed 749 metagenomic datasets from the Human Microbiome Project and identified likely previously unreported kit contamination. Collectively, our results highlight that Squeegee can identify microbial contaminants with high precision. Squeegee is open-source and available at: https://gitlab.com/treangenlab/squeegee

## Introduction

In recent years, the field of metagenomics has grown at a fast pace thanks to next-generation sequencing technologies. The scale and complexity of metagenomics studies have expanded alongside the size of the sequencing data. By performing metagenomic sequencing, we are able to analyze the DNA and RNA of the entire microbial community in varying and heterogeneous biomass environments such as samples from wastewater, soil, or human body sites^1^. One commonly used method is 16S rRNA gene sequencing. The 16S rRNA gene is highly conserved in bacteria and can be amplified and used as a marker gene for taxonomic classification^2–7^. The other widely used technique is whole-genome shotgun sequencing, where all DNA sequences in the community are fragmented and sequenced^2,3,8–10^. Both methods open the door for identifying members of microbial communities from the sampled environments and estimating the relative abundance of each member^1^. However, the results from both of these methods can be affected by microbial contamination. Microbial contamination occurs when sequences from microbes appear in the data that were not in the original samples^3,11^.

Contamination can be brought in by a variety of sources. External sources include human bodies, the laboratory environment, and kits and reagents used for collecting and processing samples^2,3,11–19^. Internal sources of contamination are often caused by mixing up different samples, such as during the sampling and sequencing process^3,11,17,20^. Contaminant sequences have also made their way into public reference databases^21–24^. Studies have shown that contaminants in DNA extraction kits are ubiquitous^25,26^, and can have critical impacts on metagenomic studies, especially for low-biomass environments, when they are not accounted for in the analysis^11^. For example, in a recent nasopharyngeal microbiota study on new born babies conducted in Thailand, contaminants found in DNA extraction kits caused an initial data analysis to become biased by the contaminants^3^.

Extra precautions during sample collection and processing, and well designed experiments, such as processing samples in a clean, well structured environment, or using depletion methods to remove host DNA, can help minimize the impact caused by contamination^11,27^. In addition, computational models have been used to identify and remove contaminants from sequenced datasets. For example, the recently published software Recentrifuge uses a score-oriented comparative approach to identify and remove contaminants from samples^28^. As is the case with all current computational methods for microbial contaminant detection, performing contamination removal with Recentrifuge requires experimental controls. Another statistical tool for identifying and removing contamination is Decontam^3^. Decontam includes a combination of a frequency-based approach and a prevalence-based approach. Auxiliary DNA quantitation data is required to perform the frequency-based analysis and standard negative control samples are required to perform the prevalence-based analysis^3^.

Experimental negative control samples combined with computational contamination identification and removal is effective^3,19,28^. However, generating experimental negative controls can be time consuming and expensive. Researchers have to perform extra experiments and do extra sequencing runs on empty samples to generate these controls. This extra work means that people must spend resources, including time and money, and as a result negative controls are often not generated. Although contaminant sequences have been a known issue for some time, negative control data are often not available in public databases, making it nearly impossible to perform contamination removal on uploaded data.

Since the composition of contaminants within DNA extraction kits and other lab reagents are ubiquitous and can be distinct, our hypothesis is that contaminants from the same sources, such as DNA extraction kits or from a lab environment, will share similar characteristics in the composition of their contaminants. This fact should enable contaminants to be found in the form of shared species in samples taken from sufficiently distinct ecological niches, or in our case, body sites. In particular, this proposed approach is most relevant when the sequencing runs use the same DNA extraction kit and/or are processed in the same lab after reaching a sufficient level of sequencing depth.

## Results

In this work, we have implemented a *de novo* computational contamination detection tool, Squeegee, which is able to identify potential contaminants at the species level. Squeegee performs taxonomic classification and searches for shared organisms across multiple samples and sample types. The workflow of the pipeline is shown in Figure 1. The software takes multiple samples containing sequencing data collected from distinct microbiomes as input, and then uses taxonomic classification to search for candidate contaminant species that are shared across samples. By estimating pair-wise similarity between metagenomic samples which the candidate contaminant species presents, and calculating breadth and depth of genome coverage by aligning the reads to the reference genome of the candidate contaminant species, we are able to identify classification errors and make accurate contaminant predictions at the species rank by filtering false calls from the candidates. We evaluate Squeegee on 3 datasets including a simulated dataset with ground truth contaminants, a real dataset with negative controls, and HMP samples without negative controls but with associated DNA extraction kit contaminants. Details on the implementation and evaluation of Squeegee can be found in methods section.

**Figure 1.**
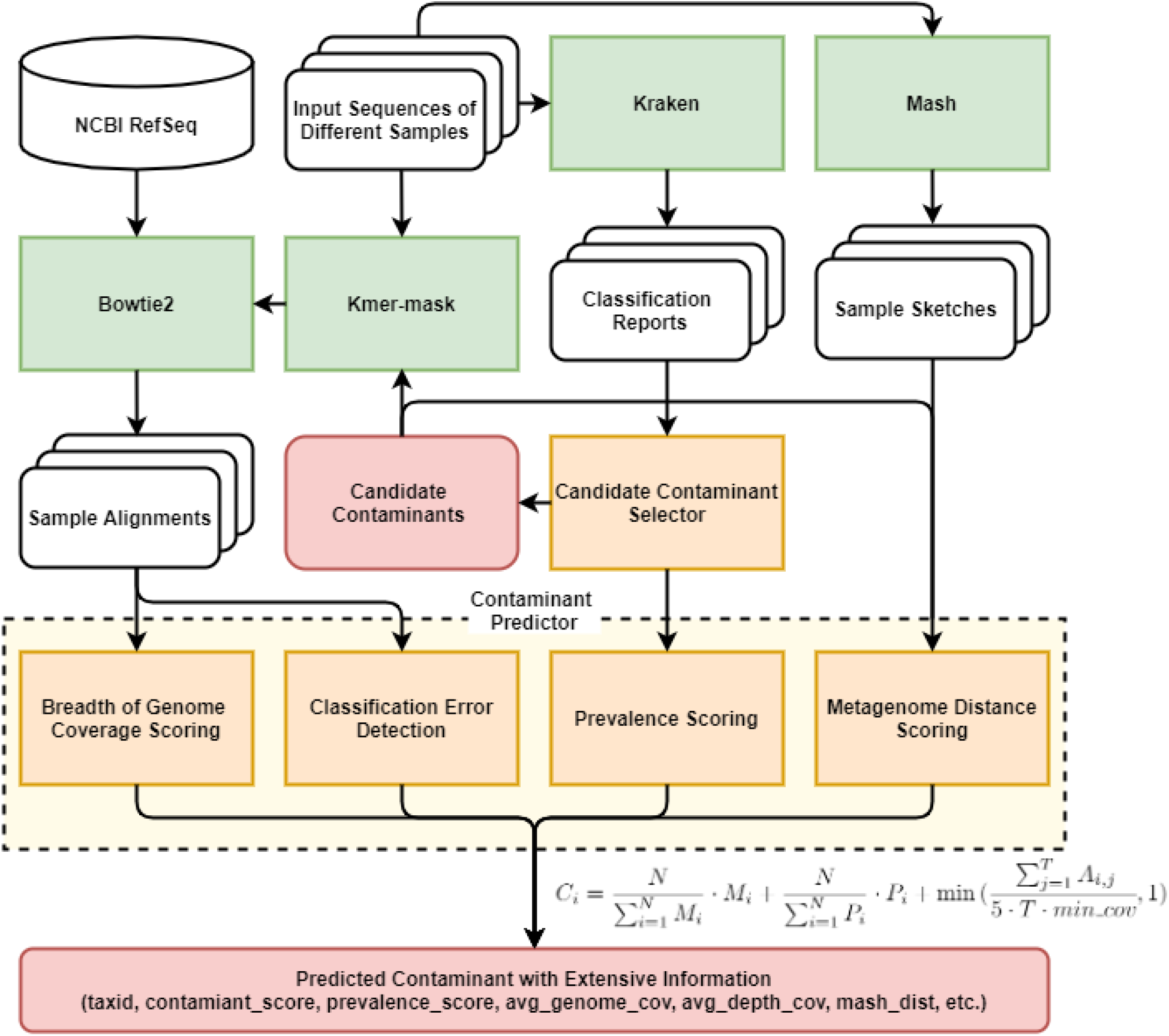
Squeegee pipeline workflow. Squeegee starts with taxonomic classification using Kraken to determine a set of candidate contaminant species. Reads from the input data are aligned to the representative genomes of the candidate contaminant species using Bowtie2 in multi-alignment mode. It also calculates the pair-wise Mash distance for all the samples. It combines the prevalence, the Mash distance, and breadth/depth of genome coverage of the candidates to predict potential contaminants.

### Stable community members for human body sites

In order to accurately identify contaminant sequences from external sources such as lab environments or reagents used during the extraction or sequencing process, the stable community members from different sample types must be considered. To assess whether there are ubiquitous genera across body sites comprising the human microbiome, we identified the stable community members across different human niches using Kraken classification results for HMP samples (Supplementary Table ST 2). By looking at each set of common community members of different body sites, we found no genera to be present in more than three of the six body sites (oral, nasal, skin, stool, throat, vaginal).

### Genus level accuracy of Squeegee prediction

We evaluated Squeegee prediction accuracy at both genus and species rank. Figure 2 shows the precision, recall, and weighted recall of Squeegee predictions in both real metagenomic datasets. For the maternal/infant dataset at genus rank, Squeegee achieved a precision of 0.833 (10/12 genera) and a recall of 0.714 (10/14 genera), where 10 correctly predicted genera occupy over 93.7% of the contaminant reads in the ground truth, indicting that Squeegee is able to identify the majority of the contaminant genera. For the HMP dataset at genus rank, Squeegee achieved a precision of 0.5 (20/40 where 20 correctly predicted genera occupy over 69.1% of the contaminant reads in the ground truth) and a recall of 0.328 (20/62 genera). Although only 20 genera were correctly predicted, the total relative abundance of those genera is over 69% of the total ground truth contaminant reads. Figure 4b shows the relative abundance of true contaminant genera identified in the MoBio DNA extraction kit. The contaminants successfully predicted by Squeegee are colored in blue and the contaminants Squeegee failed to predict are colored in red. For the simulated dataset, Squeegee achieves 100% precision and recall and is able to accurately identify all 10 spiked species among 8 different genera.

**Figure 2.**
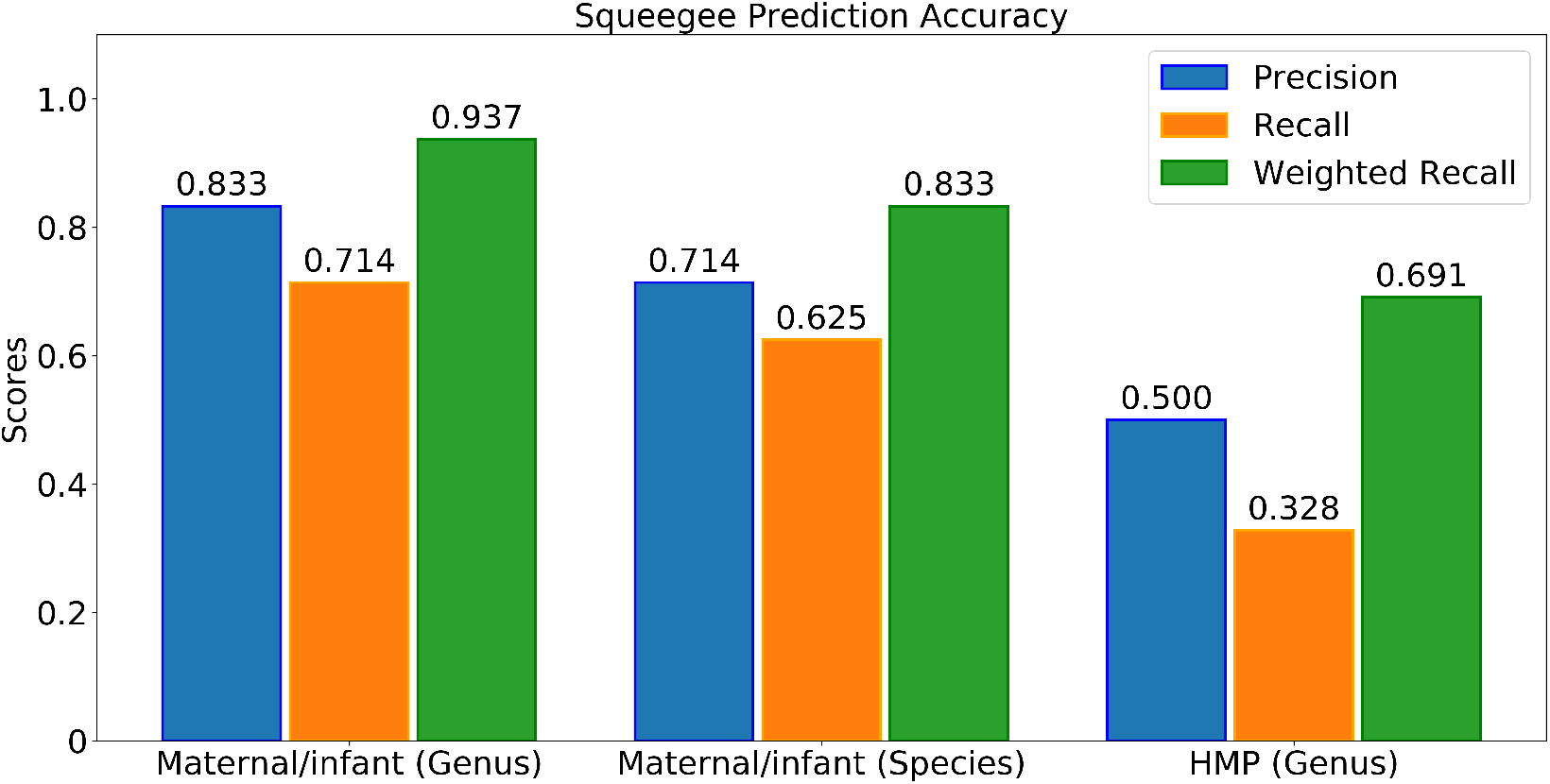
Squeegee prediction accuracy at genus and species ranks. The precision is calculated as the ratio between the number of predicted contaminant taxa found in the ground truth and the total number of predicted contaminant taxa. The recall is calculated as the ratio between the number of predicted contaminant taxa found in the ground truth and the total number of taxa in the ground truth. The weighted recall is calculated as the proportion of the reads assigned to the correct predicted taxa over the total number of reads assigned to the ground truth contaminant taxa.

### Species level accuracy of Squeegee prediction

Squeegee correctly predicted 100% of the contaminant species for the simulated dataset. For the maternal/infant dataset, Squeegee correctly predicted 10 out of 16 contaminant species observed in the contamination ground truth generated by experimental negative control samples, and achieved a recall of 0.625. The false positive calls include *Cutibacterium acnes, Rothia mucilaginosa, Staphylococcus cohnii, Staphylococcus haemolyticus*, which led to a precision of 0.714. The correctly predicted species occupy more than 83% of the relative abundance of the contamination ground truth (See Figure 2). Figure 3 shows the relative abundance of all predicted contaminant species found in each of the samples. The samples are clustered by sample type, which are shown with different colors on the color label on the y-axis. The predicted contaminant species that can be found in the negative control sample are labeled in purple at the top of the figure and the predicted contaminant species not found in the negative control sample are labeled in blue. Figure 4a shows the relative abundance of the true contaminant species identified in experimental negative control samples. The species that Squeegee successfully predicted are colored in blue and the species Squeegee failed to predict are colored in red.

**Figure 3.**
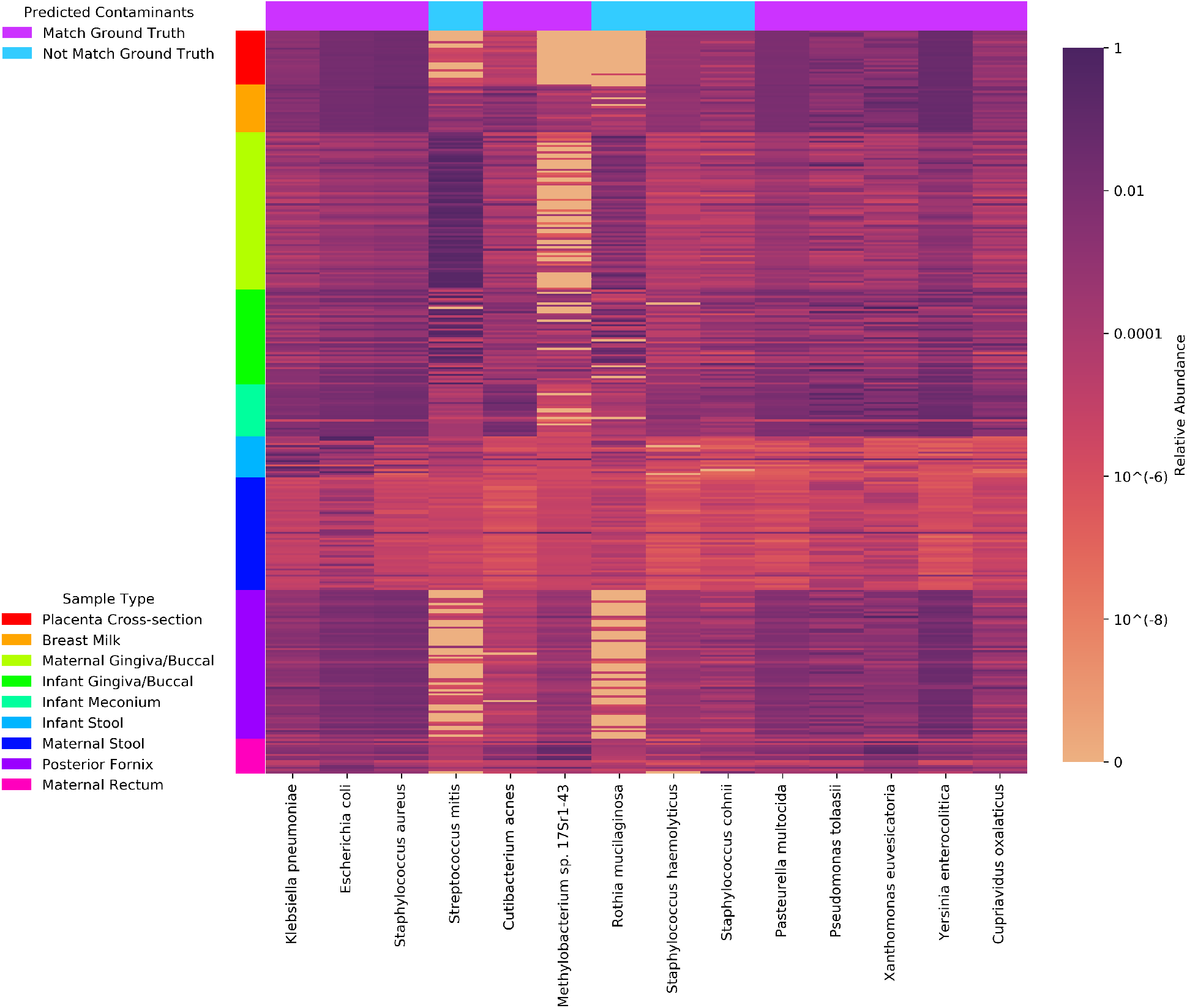
Relative abundance of all predicted species in the maternal/infant dataset. The samples are clustered by their sample type, which is shown with different colors on the color label on the y-axis. The predicted contaminant species that can be found in the negative control sample are marked by the purple label on the x-axis, whereas the predicted contaminant species that cannot be found in the negative control sample are marked in blue.

**Figure 4.**
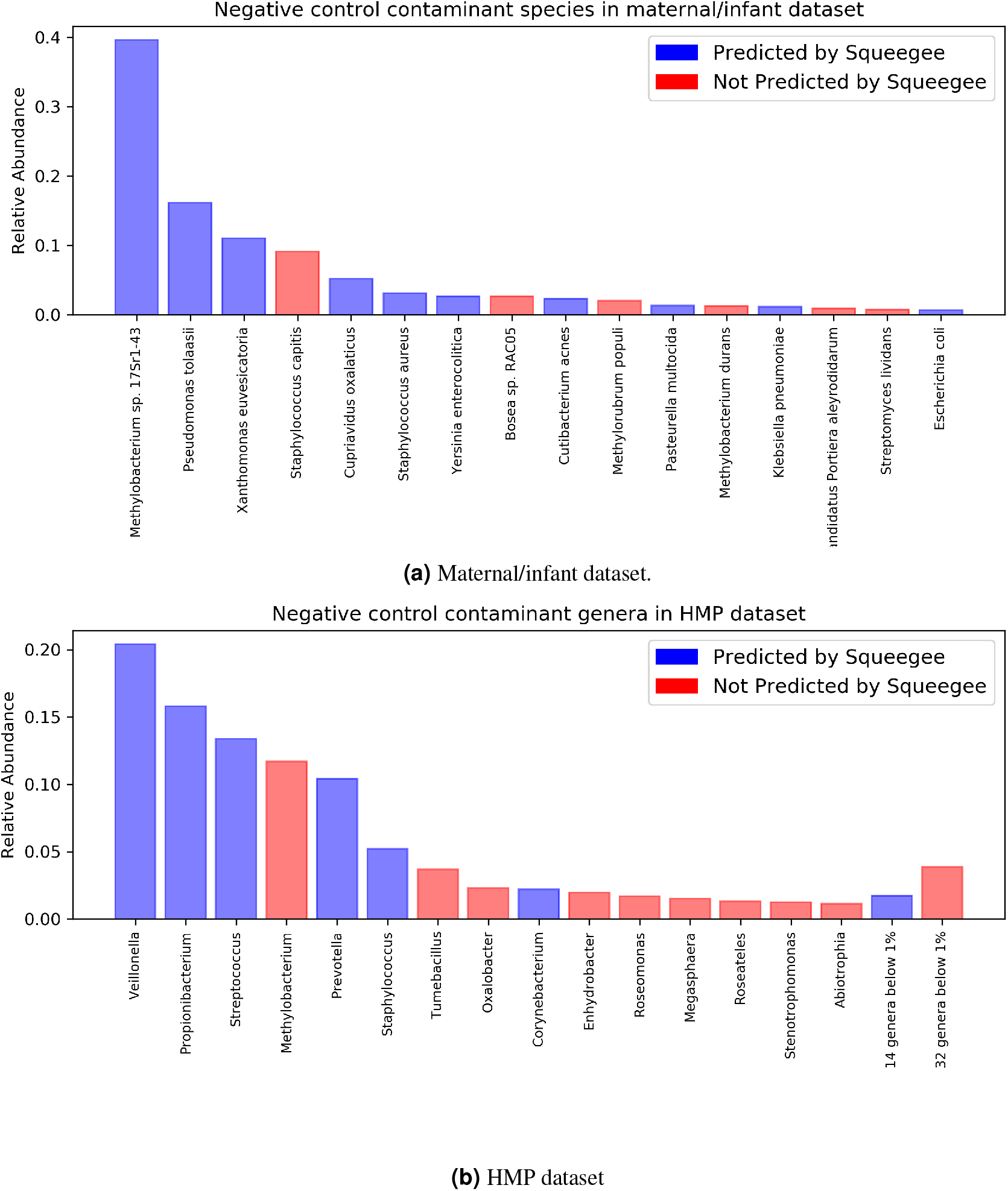
Relative abundance of ground truth contaminants of (a) maternal/infant and (b) HMP dataset. The correct predicted species/genera are marked in blue, and the species/genera that Squeegee failed to predict is marked in red. For HMP dataset, genera with relative abundance below 1% are combined.

For the HMP dataset, since we are using bacteria identified at the genera level as inherent contaminants in the MoBio DNA extraction kit level^18^ for our negative control reference, recall and weighted recall calculations at the species level do not apply. Thus, the precision is calculated as the number of predicted contaminant species that fall under the negative control genus out of the total number of predicted contaminant species. Squeegee achieved a precision of 0.762 (92/126 species). Figure 5 shows the prevalence, the breadth of genome coverage, and additional score and filtering information of the top 50 predicted contaminant species after filtering. The first 16 rows show the prevalence of each species among each of the sample types, where zero prevalence is marked in blue. The next 16 rows show the breadth of genome coverage of each species in each of the sample types. The remaining rows show the prevalence score, the alignment score, the Mash score, and the combined score used to make the final prediction, and whether each species passes the filters. The last row of the heat map shows whether the species can be found in the ground truth, with true positives shown in white and false positives shown in black. Detailed information of all candidate contaminant species can be found in Supplementary Fig. 1.

**Figure 5.**
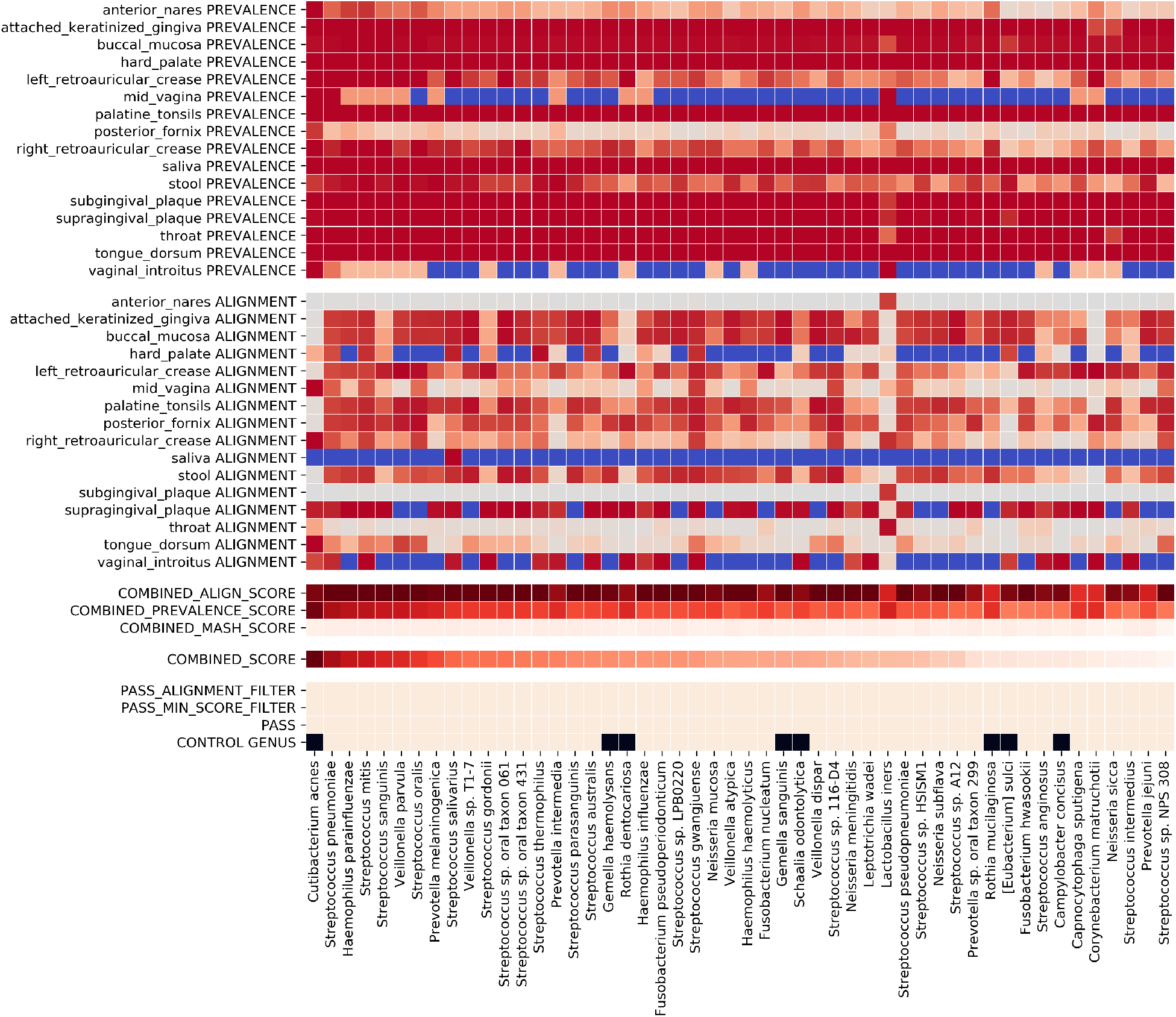
Scoring and filtering of candidate contaminants for the HMP dataset. This plot shows the prevalence, the breadth of genome coverage, and additional score and filtering information of the top 50 contaminant species after filtering. The first 16 rows show the prevalence of each species among each of the sample types, where zero prevalence is marked in blue. The next 16 rows show the breadth of genome coverage of each species in each of the sample types. The remaining rows show the prevalence score, the alignment score, the Mash score, and the combined score used to make the final prediction, and whether each species passes the filters. The last row of the heat map shows whether the species can be found in the ground truth with true positive show in white and false positive show in black.

### Alpha diversity analysis before and after contamination removal

Figure 6a shows Shannon’s diversity index and Simpson’s diversity index for the maternal/infant dataset before and after contamination removal. Both diversity metrics for the samples were evaluated before the contaminant reads were removed (shown in red), after removing species confirmed by the experimental negative control (shown in blue), and after removing all species predicted by Squeegee (shown in black). The max removal cutoff is set to 1%, which only removes species with relative abundance less than 1%. We observed significant decreases of Simpson’s diversity index in both placental and breast milk groups and significant decreases of Shannon’s diversity index in the placental group. There are also significant decreases of Shannon’s diversity index in the breast milk group if we remove all predicted contaminant species, but no significant decreases are found by only removing contaminant species confirmed by the negative control experiments. For a more strict max removal cutoff of 0.5%, we still found significant decreases of both Shannon’s and Simpson’s diversity index in the placental group (See Supplementary Fig. 2).

**Figure 6.**
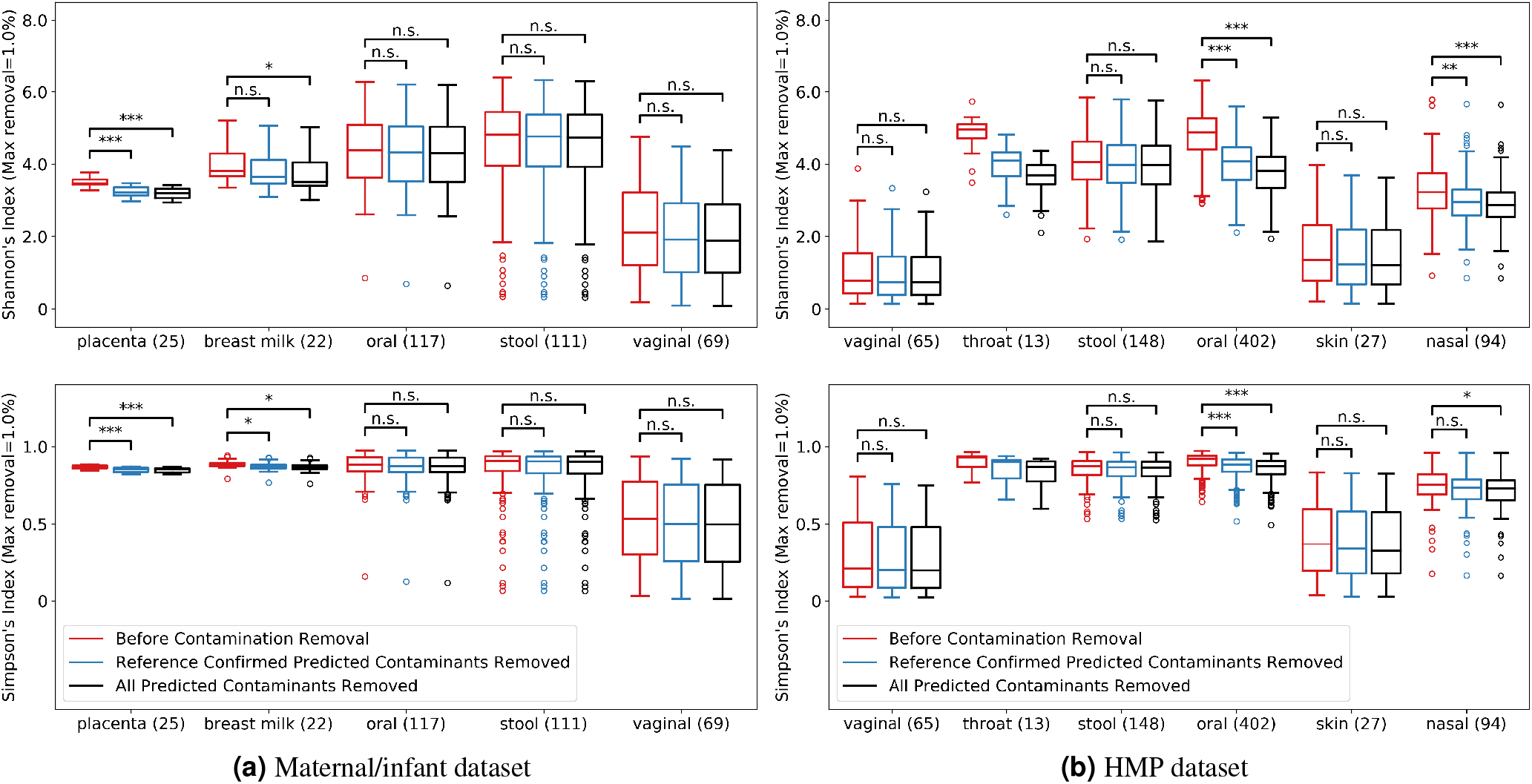
Alpha diversity index for (a) maternal/infant dataset and (b) HMP dataset. Both Shannon’s and Simpson’s diversity index of the communities in each of the samples were evaluated before the contaminant reads were removed (red), after removing species only confirmed by the experimental negative control (blue), and after removing all species predicted by Squeegee (black). The max removal is set to 1%. Numbers inside parentheses are the numbers of samples in each sample type. Significance labeling: n.s.(P*>*0.05), *(P*≤*0.05), **(P*≤*0.01), ***(P*≤*0.001).

Figure 6b shows the same alpha diversity analyses performed on the HMP samples with the maximum removal cutoff set to 1%. We observed significant decreases for Shannon’s diversity index values in oral and nasal samples, and a significant decrease in Simpson’s diversity index in oral samples. Significant decreases of Simpson’s diversity index are found in the case of removing all predicted contaminant species, but there are no significant decreases in the case of only removing contaminant species confirmed by the MoBio contaminants. With the max removal cutoff of 0.5%, there are significant decreases of Shannon’s diversity index in the oral and nasal-pharyngeal samples, and Simpson’s diversity index in the oral samples (See Supplementary Fig. 3).

## Discussion

Squeegee is the first *de novo* computational tool designed to identify and mark suspicious taxa as potential contaminants in the absence of “kit negative” or environmental contaminant controls. Squeegee is able to mark these taxa contained within metagenomic samples without requiring negative experimental controls. In order to predict contaminant species, multiple pieces of evidence are taken into consideration, including the prevalence rate of species, the metagenomic distance of the samples that contain the species, and how well the genomes of those species are being covered. Comparisons between Squeegee predictions and experimental control data show that Squeegee is capable of accurately inferring contamination at the species level, especially in regards to contaminants occurring at a high relative abundance.

In the maternal/infant dataset, we observed that Squeegee only failed to predict a few contaminant species found in the negative control samples. Those false negative contaminants all have relative abundances below 5% except for *Staphylococcus capitis*. We also found that Squeegee predicted a number of species from the genera *Staphylococcus* including *Staphylococcus haemolyticus, Staphylococcus mitis, Staphylococcus cohnii*, that are not found in the experimental control samples. The only other false positive call was *Rothia mucilaginosa. Staphylococcus* species are often found in the normal flora of the skin, and have been reported multiple times as contaminants from DNA extraction kits and laboratory environments^18,29^. *Rothia mucilaginosa* is a part of the normal oropharyngeal flora and has also been found in DNA extraction kits^18,30^. It is possible that the experimental control samples were not sequenced deeply enough to reveal these species, or the species were at a low enough relative abundance in the experimental control samples that they were filtered out during quality control.

Squeegee is designed for de novo identification of microbial species that are likely contaminants; a higher combined contaminant score indicates the species has a higher potential for being an actual contaminant. However, Squeegee failing to flag a microbial species in a sample as a likely contaminant does not mean it is not a contaminant. One of the limiting factors is the relative abundance of the species within the source of the contamination. Figure 4a and figure 4b show that contaminant species with low relative abundances in the control samples are more difficult to identify, since the sequencing signals of such species become even weaker in the non-control metagenomic samples. One of the other limitations of Squeegee is that it cannot trace contaminants originating from the sample collection process, since different sample collection operations may introduce different contaminant species. Therefore, for species that are not included in the predicted contaminants, further investigation is required to validate whether the species truly originated from the sampled metagenome. Squeegee can help rule out misclassification,

Since Squeegee operates without prior knowledge of the input dataset, ubiquitous species that are commonly found in a wide range of environments could allow Squeegee to make false predictions. Although *Staphylococcus* genera have been reported as external contamination from multiple studies, it is hard to ignore the fact that some of the *Staphylococcus* species may be truly present among multiple different body sites, including skin and nasal samples. Such ubiquitous species may introduce noise in Squeegee’s predictions. Combined with the prior knowledge of the input dataset and the comprehensive information that Squeegee outputs, the user may further filter the predicted list of contaminants if needed.

By no means is Squeegee a replacement for experimental negative controls, and it does not estimate relative abundance of each predicted potential contaminant since the relative abundance of the contaminants varies in different sample types. Squeegee makes predictions based on the assumption that the input data are sampled from multiple distinct microbiomes, and does not apply to cases where the sequencing data are from similar microbiomes. If possible, performing negative control experiments will provide a more accurate profile of the external contaminants. But, as discussed, it is not uncommon that experimental negative control samples are not available for a huge number of public datasets. The data from the Human Microbiome Project is one high profile example of this. When compared to other contamination removal methods, Squeegee is the only existing tool able to predict contamination from multiple sources without experimental negative control samples (see table 1), and its contaminant predictions can have a significant impact on diversity measures which are often a key part of the results of a vast range of microbial studies^3,22,28,31,32^.

**Table 1.**
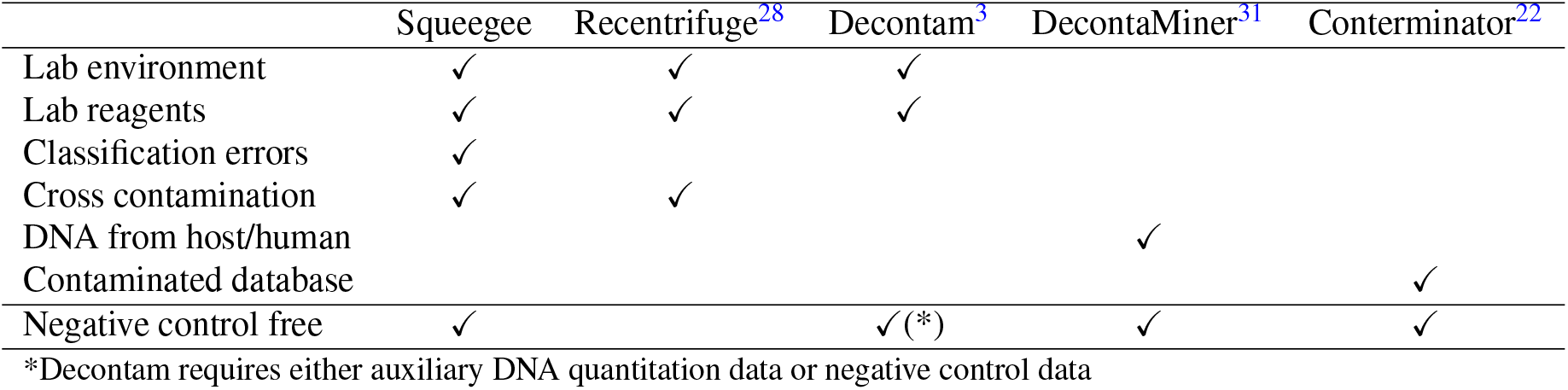
Tools comparison on handling contamination from different source

Another possible solution for contamination detection without negative control sequencing data is to use a contaminant database. If there exists a database of genomes containing known contaminant species, we could identify the contaminant sequences in the data by mapping reads against this database^33^. Building such a contaminant database can be challenging because it requires sequencing data from all possible sources of contamination. Since Squeegee is a negative control free tool for identifying novel contaminants, it can also used as an important step in filling out such a comprehensive database of putative likely contaminants.

Over 76% of the contaminant species predicted by Squeegee for the HMP dataset match the bacterial genus described as inherent contaminants of the MoBio DNA extraction kit, which was used for the Human Microbiome Project^18^. The Squeegee prediction has a weighted recall of 0.69, and Squeegee failed to predict most of the genera from phylum *Proteobacteria*. This may be due to the fact that the kit used in the Mobio contamination study^18^ is close related to the one used for HMP, but not identical. The contamination profile of the same kit might change over time, and samples processed in different labs may also affect the results since contaminants from lab surfaces and lab members have the potential to contribute to the composition of the contamination.

It is worth pointing out that a stable community member of a certain body site has the potential to also be a contaminant taxon from an external source. For example, species from the genera *Staphylococcus* are commonly found in skin samples, but they are also commonly found as an inherent contaminant in multiple DNA extraction kits. In general, Squeegee makes contamination predictions based on shared species across different sample types. For any individual sample type, the user should treat the predicted result with care to avoid potential community members falsely labeled as contaminants.

Finally, Squeegee was tested and evaluated with metagenomic shotgun sequencing datasets. Under the same hypothesis, Squeegee could be readily altered and extended for use on 16S rRNA sequencing data. In such a case, we are not be able to use breadth and depth of genome coverage of the alignment to determine classification errors. Therefore, choosing an accurate taxonomic classifier is critical for running Squeegee on 16S rRNA sequencing data.

In summary, Squeegee is the first computational method for identifying potential microbial contaminants in the absence of environmental negative control samples. Squeegee predictions on multiple datasets have shown that contaminant sequences from the same source, such as DNA extraction kits and other reagents used during the sample processing and sequencing, can be accurately identified across multiple samples using this computational method without experimental negative controls. Squeegee achieves both high weighted recall and low false positive rates on real metagenomic datasets, and can help to identify putative contaminant sequences of suspicious taxa for low biomass microbiome studies, enabling sample-independent and orthogonal approaches aimed at distinguishing true microbiome signals from environmental contamination.

## Methods

### Samples from distinct environments

In order to generate reproducible estimates of contaminants and their composition among the samples, the user must collect sequencing data from multiple metagenomic samples. The microbial community composition should be largely distinct between any two samples included in the analyses. Here, distinct refers to different metagenomic environments or sample types in which it is rare to observe a given microbial species present across most samples. Each sample should be provided with a tag or descriptor that distinguishes the different types of samples (*e*.*g*. oral, vaginal, fecal, soil, ocean, etc).

### Taxonomic classification

Squeegee first performs taxonomic classification using Kraken v1.1.1^34^ with default settings (k=31). The reference database for Kraken was build with complete bacterial/archaeal/viral genomes from NCBI RefSeq (Release 202). A classification report is generated for each of the samples. Based on the classification, Squeegee chooses a set of candidate contaminant species based on the prevalence of the species across the samples. The prevalence score is weighted by the number of samples of the same type to avoid bias introduced by an unbalanced number of samples between sample types. Higher prevalence rates of a species indicates that the species is shared by more samples across more sample types, and it is more likely to be a contaminant.

### Metagenomic distance estimation

Squeegee also calculates the metagenomic similarity between the samples using Mash v2.2.2, a tool which estimates the Jaccard index using MinHash^35^. This is done by first generating a sketch of each sample (Mash sketch -s 100000 -k 21 -m 2) and then calculating the pair-wise Mash distance between all pairs of samples (Mash dist). High Mash distances indicate the metagenomes of two samples are more distinct (i.e. there are fewer genera and species shared between the samples). Squeegee weights shared species coming from more distinct samples as more likely to be a contaminant.

### Read alignment and error identification

Squeegee then fetches the representative genomes for each of the candidate contaminant species from the NCBI RefSeq database used to build the Kraken database. These representative genomes are used as references to perform a multi-alignment for all reads in the samples using Bowtie2 v2.3.5 with the multi-alignments enabled (bowtie2 –local -a –maxins 600)^36^. To accelerate this process, kmer-mask from meryl v1.0 is used to filter out reads that do not contain any 28-mers from the reference genomes (kmer-mask -ms 28 -clean 0.0 -match 0.01 -nomasking)^37^. Based on the alignment results, the breadth and depth of genome coverage is calculated for each of the sample types using samtools v1.11 (samtools depth)^38^. The breadth and depth of genome coverage is used to determine whether the species is truly present or if the species is a potential misclassification from the taxonomic classifier. A species that is truly present should have a large proportion of its genome covered. On the other hand, a large number of reads covering only a small proportion of the genome often suggests that the species was a misclassification^33^. Since contaminant species are often low in abundance, combining samples from the same type would give us a better indication of the presence of the species.

### Contaminant predictions

In the last step, Squeegee combines multiple pieces of evidence including the prevalence score, Mash distance score, and alignment score and makes a final prediction for contaminant species using equation 1,

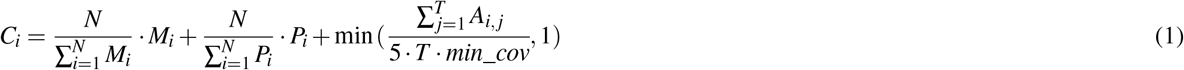

where *N* is the total number of candidates, and *T* is the total number of sample types. *M*_*i*_ is the Mash distance score of candidate contaminant species *i*. We took the Mash distance values (from 0 to 1) of all sample pairs that both contain species *i*, and calculate *M*_*i*_ by averaging the top 10% of the pairwise Mash distance value. We defined *P*_*i*_ as the prevalence score of candidate contaminant species *i*, which is calculated as the mean prevalence rate of species *i* among all sample types. *A*_*i, j*_ is the alignment score of candidate contaminant species *i* in sample type *j*, which is defined as the breadth of genome coverage of species *i* in sample type *j* with minimum depth of 3.

After the combined contaminant scores are calculated, Squeegee filters out species that are below a user defined minimum combined score threshold. Candidate contaminants with a low combined score suggest that there is not enough evidence supporting the argument that the candidate species is both a true contaminant and definitely present in the samples. Squeegee also provides a comprehensive output for the user if further downstream analysis is required.

### Evaluation of Squeegee

Evaluation of Squeegee predictions was performed by comparing the predicted contaminant species using three datasets: (1) a simulated dataset with ground truth contaminant species, (2) a real dataset with available negative control samples, and (3) a real dataset without a negative control (HMP samples) but with associated kit contaminants. For (1), the simulated dataset, the contaminant species in the ground truth were generated based on the species of a simulated spike-in of contaminant sequences. A total number of 18 simulated samples were generated using CAMISIM and ART simulating Illumina paired-end reads with average read length of 150bp^39,40^. The total number of read pairs in each of the simulated samples is 3322898, containing true sample species sequences and spiked-in contaminant sequences. All simulated samples are divided into 6 groups, 3 samples per group. Each group of samples contained sequences from 5 different bacteria genomes which serve as true organisms in the sampled community (distinct among groups), and sequences from 10 common contaminant bacteria genomes (shared among groups). The relative abundances of spiked-in contaminant sequences are 0.05, 0.10, and 0.20 for three simulated samples in each group. For (2), maternal/infant metagenomic datasets, the contaminant species in the ground truth was generated based on the classification of multiple experimental negative controls. To minimize classification errors, we applied a set of criterion to include a species in the contamination ground truth. Species with relative abundance above 0.5% or more than 3000 reads assigned in more than half of the negative control samples, and species with relative abundance above 10% in a single sample were chosen for inclusion in the ground truth contaminant set. We then aligned the sequencing reads in the experimental control samples to the representative genomes. Reads assigned to the *Staphylococcus virus Andhra* stacked in a small 449 bp region with an average depth of 1429, indicating a false classification call, so we removed it from the ground truth contaminants. Once the ground truth contaminants were identified, the relative abundance of the ground truth contaminants is calculated as the average relative abundance across all negative control samples over the sum of the average relative abundance of each contaminant. For (3) the HMP dataset, which was extracted using the MoBio DNA extraction kit, we used the 62 bacteria (excluding lot dependent organisms) which were identified as inherent contaminants within a latter version of a related MoBio extraction kit, the MoBio PowerMax_®_ Soil DNA Isolation Kit 12,988-10 (MoBio Laboratories, USA), in a recent study^18^ as the ground truth contaminants. Relative abundances of each genera were also obtained in the same study. Since Squeegee makes contamination prediction at the species level, predicted contaminant species from the reference genus are counted as true positives.

The accuracy of the prediction is measured by precision, recall, and weighted recall. The precision is calculated as the ratio between the number of predicted contaminants found in the ground truth and the total number of predicted contaminants. The recall is calculated as the ratio between the number of correctly predicted contaminants and the total number of contaminants in the ground truth. The weighted recall is calculated as the proportion of the reads assigned to the correctly predicted contaminants over the total number of reads assigned to the ground truth contaminants. Accuracy at the genus level is calculated using the genera of each predicted species as the predicted contaminant genera. The parameters and data characteristics are shown in Supplementary table 1.

The simulated datasets is publicly available and can be downloaded at https://rice.box.com/s/x3645qvtswvb838e0dfsvk0gre74j9gz. The maternal/infant metagenomic datasets are available for download via NCBI BioProject PRJNA725597. The HMP samples are downloaded from https://www.hmpdacc.org/HMASM/. The datasets were simulated/sequenced from distinct samples, and no sample was simulated/sequenced repeatedly.

### Alpha diversity analysis of predicted contaminants

We categorized the labeled sample types of the maternal/infant data set and HMP data set into combined sample types based on body site. The combined sample types for the maternal/infant data set include placenta, breast milk, oral, stool, and vaginal. The combined sample types for HMP includes vaginal, throat, stool, oral, skin, and nasal samples. Samples from the same combined sample types in each data set were used for alpha diversity analysis. Both Shannon’s diversity index and Simpson’s diversity index were measured before and after contamination removal. Only reads assigned to the species rank by Kraken were used in calculating Shannon’s diversity index and Simpson’s diversity index. Since contamination originating from external sources can also be true community members of the metagenomes, we also set a max removal cutoff and only remove species with relative abundance below this cutoff. The significance test was done using two-sided Mann–Whitney *U* test for all combined sample types with more than 20 samples.

### Stable community members for human body sites

We used the samples from the HMP data set and their combined sample types to generate a set of stable community members for different human body sites. Stable community members were defined as genera with more than 1% of their reads assigned from Kraken classification in more than 50% of the samples from the same combined sample types.

## Acknowledgements

We would like to thank Michael Nute for constructive comments and helpful feedback. We would also like to thank Ben Callahan for his feedback on Decontam algorithm. T.T. was supported in part by NIH grant 1P01AI152999-01 supported by National Institute of Allergy and Infectious Diseases (NIAID) L.E, Y.L, and T.T. were supported by the FunGCAT program from the Office of the Director of National Intelligence (ODNI), Intelligence Advanced Research Projects Activity (IARPA), via the Army Research Office (ARO) under Federal Award No. W911NF-17-2-0089. This research was made possible in part by an NIH-funded fellowship to Dr. Michael Jochum (T32 HD098069).

## Author contributions statement

L.E., K.A, T.T, conceived the experiment(s), Y.L. conducted the experiment(s), M.J, L.E, K. A., T.T. and Y.L. analysed the results. All authors reviewed the manuscript.

