## SUPPLEMENTARY DATA for "Squeegee: de-novo identification of reagent and laboratory induced microbial contaminants in low biomass microbiomes"

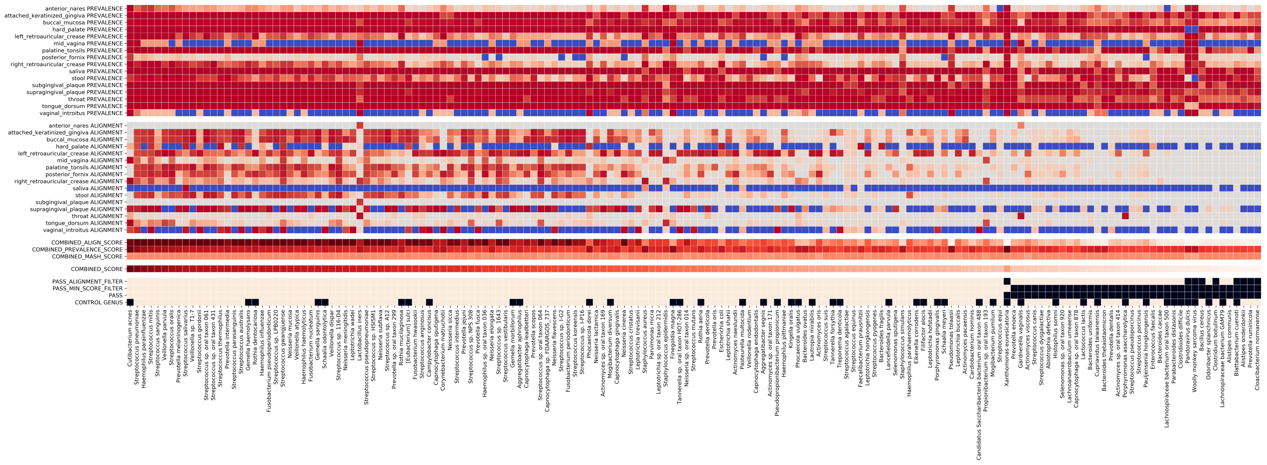


**Supplementary Figure 1.** Scoring and filtering of candidate contaminants for the HMP dataset. This plot shows the prevalence, the breadth of genome coverage, and additional score and filtering information of candidate predicted contaminant species. The first 16 row shows the prevalence of each species among each of the sample type, where zero prevalence is marked in blue. The next 16 rows shows the breadth of genome coverage of each species in each of the sample type. The remaining rows shows the prevalence score, the alignment score, the Mash score, and the combined score used to make the final prediction, and whether each species passes the filters. The last row of the heat map shows whether the species can be found in the ground truth with true positive show in white and false positive show in black.

**Supplementary Table 1.** Parameters and Dataset Characteristics

| | Simulated | maternal/infant | HMP |
|---|---|---|---|
| Total number of samples | 18 | 344 | 749 |
| Total number of sample types | 6 | 9 | 16 |
| Prevalence min read | 30 | 30 | 30 |
| Prevalence min abundance | 0.05% | 0.05% | 0.05% |
| Min genome coverage | 2.5% | 2.5% | 7.5% |
| Min combined score | 0.70 | 0.75 | 0.75 |
| # of ground truth species | 10 | 16 | N/A |
| # of ground truth genus | 8 | 14 | 62 |
| # of predicted species | 10 | 14 | 126 |
| # of predicted genus | 8 | 12 | 40 |
| # of correct predicted species | 10 | 10 | 92 |
| # of correct predicted genus | 8 | 10 | 20 |

**Supplementary Table 2.** Genera presence/absence across different body sites.

| | Vaginal | Throat | Stool | Oral | Skin | Nasal |
|---|---|---|---|---|---|---|
| Alistipes | | | ✓ | | | |
| Bacteroides | | | ✓ | | | |
| Campylobacter | | ✓ | | | | |
| Corynebacterium | | | | | | ✓ |
| Cutibacterium | | | | | ✓ | ✓ |
| Fusobacterium | | ✓ | | ✓ | | |
| Haemophilus | | ✓ | | ✓ | | |
| Lactobacillus | ✓ | | | | | |
| Neisseria | | ✓ | | ✓ | | |
| Prevotella | | ✓ | | ✓ | | |
| Parabacteroides | | | ✓ | | | |
| Phocaeicola | | | ✓ | | | |
| Rothia | | | | ✓ | | |
| Staphylococcus | | | | | ✓ | ✓ |
| Streptococcus | | ✓ | | ✓ | | |
| Veillonella | | ✓ | | ✓ | | |
| Xanthomonas | ✓ | | | | ✓ | ✓ |

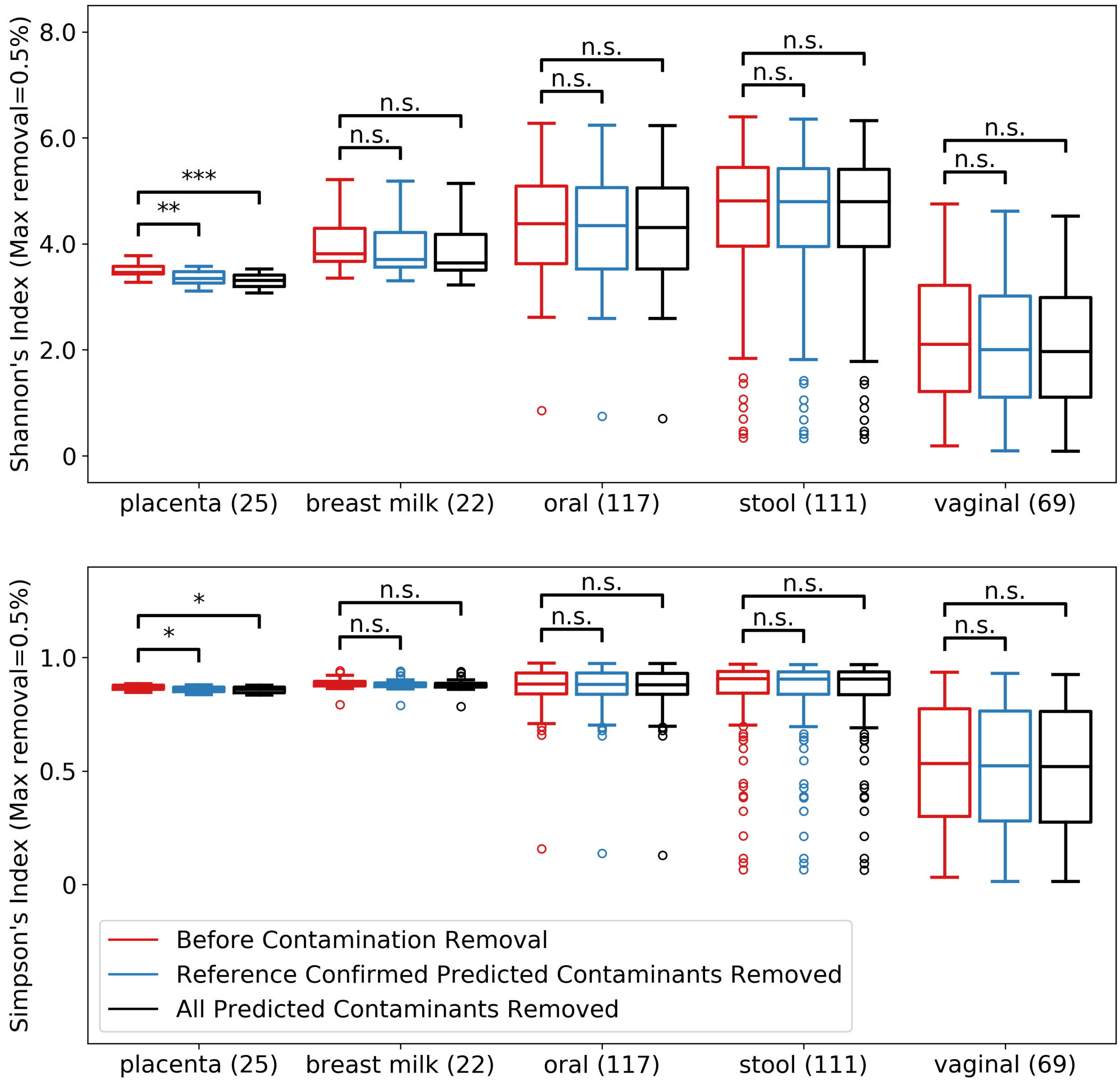


**Supplementary Figure 2.** Alpha diversity index for maternal/infant dataset. Both Shannon's and Simpson's diversity index of the communities in each of the samples were evaluated before the contaminant reads were removed (red), after removing species only confirmed by the experimental negative control (blue), and after removing all species predicted by Squeegee (black). The max removal is set to 0.5%. Numbers inside parentheses are the numbers of samples in each sample type. Significance labeling: n.s.(P>0.05), *(P≤0.05), **(P≤0.01), ***(P≤0.001).

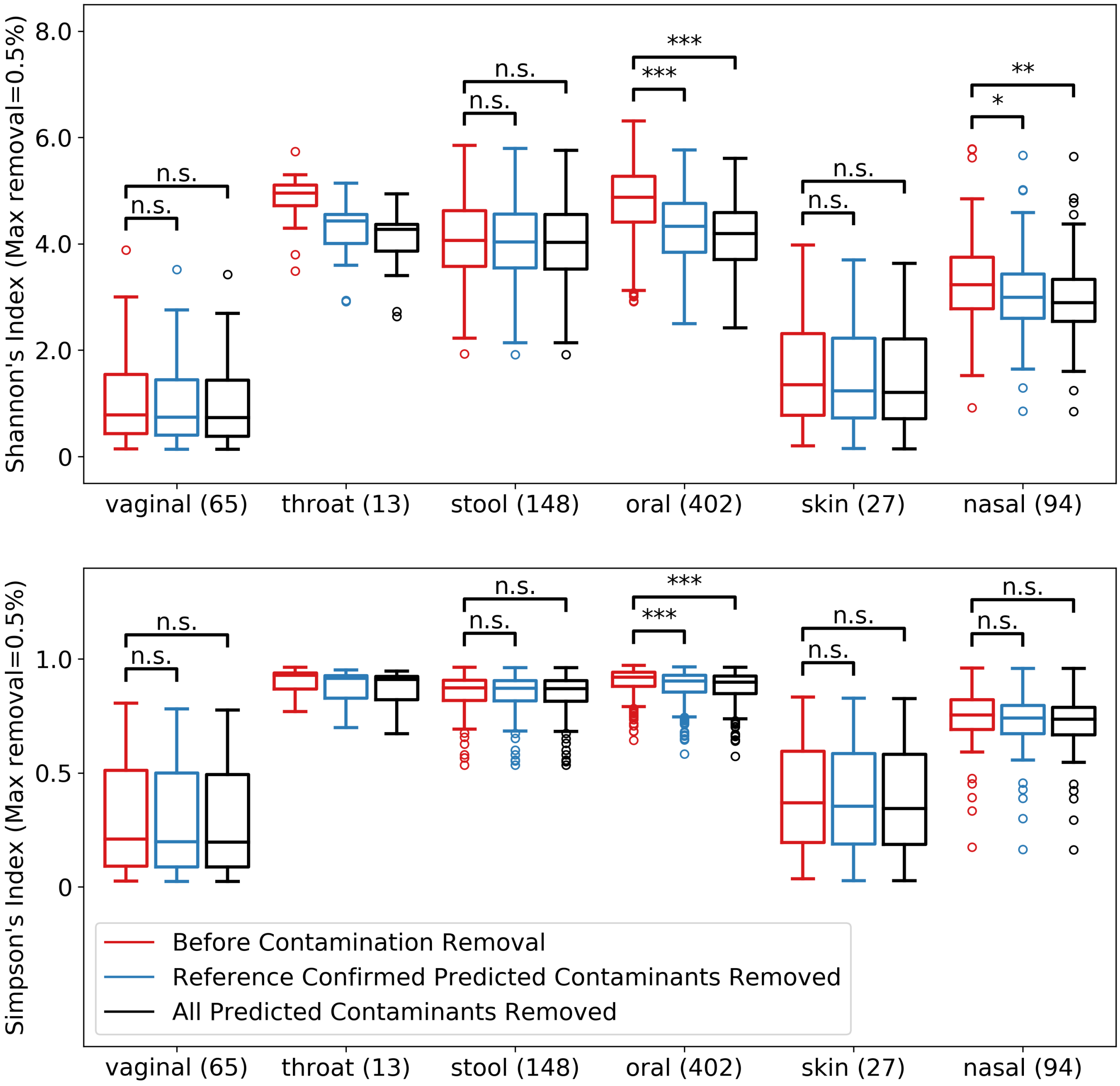


**Supplementary Figure 3.** Alpha diversity index for HMP dataset. Both Shannon's and Simpson's diversity index of the communities in each of the samples were evaluated before the contaminant reads were removed (red), after removing species only confirmed by the experimental negative control (blue), and after removing all species predicted by Squeegee (black). The max removal is set to 0.5%. Numbers inside parentheses are the numbers of samples in each sample type. Significance labeling: n.s.(P>0.05), *(P≤0.05), **(P≤0.01), ***(P≤0.001).
